# Neurodynamics of prefrontal areas in volitional contexts- a comparative study based on computational modelling and EEG/ERP data

**DOI:** 10.1101/2025.11.28.691148

**Authors:** Azadeh Hassannejad Nazir, Tomas Watanabe, Mikael Lundqvist, Nima Khalighinejad, Jeanette Hellgren Kotaleski, Hans Liljenström

## Abstract

Volitional action arises from the interaction between internal intentions and external stimuli, yet the neural dynamics distinguishing self-initiated from externally triggered actions remain unclear. Here, we combine human EEG/ERP data with simulations from a phenomenological neurocomputational model of prefrontal control to examine the neurodynamic principles underlying different types of volitional behaviors. Using an extended version of our previously developed model, we generated neural activity patterns corresponding to self-initiated and externally triggered actions by manipulating the balance between endogenous and exogenous inputs to lateral prefrontal subregions. We then qualitatively compared these simulated dynamics with empirical EEG data from a perceptual decision-making task involving voluntary and instructed skip actions. Across both datasets, self-initiated actions showed a gradual buildup of activity, marked reductions in cross-trial variability, sharper state transitions, and beta–gamma frequency shifts; in contrast, externally triggered actions exhibited minimal variability reduction, weaker transitions, and largely stable frequency content. These convergent results suggest that self-initiated actions are supported by distinct preparatory dynamics, characterized by internal competition and stabilization of neural states, whereas externally triggered actions rely primarily on externally driven input. Together, these findings identify candidate neurodynamic signatures of intention formation and highlight how simplified computational models can reproduce and help explain key features of volitional control observed in human electrophysiology.

## Introduction

Every day, we experience different feelings about our agency in performing various actions. The sense of control over our actions and thoughts, as well as the feeling of being controlled by the environment and society, represents two sides of every decision in our lives. Feelings of agency and control arise when our behaviors are consciously initiated and occur without immediate external triggers. Considering this perceived phenomenon in self-initiated volitional actions^1^, it might be possible to regard certain non-immediate, externally driven actions as similar to self-initiated actions, in that they are also volitional or freely chosen. One clear distinction between these two types of actions lies in the origin of the stimulus that drives behavior—either internally generated (endogenous) or triggered by external factors. This raises a key question: are externally triggered actions based on similar neural mechanisms and do they exhibit similar emergent neurodynamical behavior as self-initiated actions? To address this question, it is essential to consider different cognitive functions—such as perception, learning, and intentional processes—not as isolated aspects, but as hierarchically interconnected building blocks of volitional action^2^. According to Freeman^3^, understanding a perception-action cycle embedded in every human mind plays a pivotal role in exploring individual behaviors. In this dynamic process, consciousness, attention, and intention are crucial in driving the cycle from perception to action^4,5^. Therefore, investigating the preparatory process of intention and attentional control may lead us a better understanding of human behavior in a volitional context.

### Voluntary action

All through history, the significance of agency in shaping human existence has continuously sparked debate and inquiry. Addressing questions such as whether humans possess the freedom to make decisions amidst social and environmental influences lies at the heart of research on volitional control, arbitrariness, and free will across various scientific disciplines.

Understanding the volitional decision-making process is a multidimensional endeavor. In particular, the interplay among contextual factors, internal signals, actions, and their outcomes constitutes a central component of voluntary actions^6^. Within this framework, the preparatory process of intention encompasses both intentional goal control and intentional action control. More specifically, volitional decision-making involves: 1) goal-directed actions shaped by motivations that may be independent of immediate external stimuli^7^; and 2) the control of actions and the prediction of their associated outcomes; 3) an understanding of the relationship between reasons and actions. A key element in this process is the presence of consciousness, which underlies the sense of agency and provides the predictive basis for potential actions through its association with causal processes^7,8,9^.

This cognitive perspective aligns with accounts emphasizing the need for coordination across multiple brain areas during volitional decision-making. In particular, volitional behaviors, as manifestations of higher-order cognitive processes, span prefrontal and premotor cortical areas such as the lateral prefrontal cortex (LPFC) and the pre-supplementary motor area (pre-SMA), whose anatomical and functional properties suggest a crucial role in preparatory processes^10,11^.

Experimental paradigms such as the Libet task and intentional inhibition tasks have allowed researchers to observe the neural precursors of voluntary action. The most prominent marker is the readiness potential (RP), an average drift in cortical electrical activity that typically emerges 1 to 2 seconds before movement onset, peaking just prior to action initiation. The RP is thought to originate from the pre-SMA. Libet and colleagues^12,13^ demonstrated that the RP could be detected before subjects reported a conscious decision to act, sparking debate on the neurophysiological basis of free will and highlighting methodological challenges in studying the neuroscience of consciousness^13, 14^. By linking cognitive constructs of agency and causality to measurable brain activity, these paradigms provide a bridge between theory and physiology.

Dynamical systems theory offers a complementary framework to integrate these findings. This theory provides tools to analyze the brain’s intrinsic nonlinear dynamics, which manifest as high-dimensional, potentially chaotic neural patterns^15,16^. Viewing the brain as a dynamical system provides a basis for understanding the adaptation and goal-directedness of human actions. A system that remains in a steady state of unchanging equilibrium would not be capable of decision-making or adaptation to environmental or internal changes^17^, whereas a system capable of dynamical transitions provides a natural framework to associate decisions, choices between divergent or competing alternatives, and complex behavioral patterns observed across multiple timescales. This perspective leads to the hypothesis that a perception-action cycle not only correlates with but also evolves along a chaos-to-stability pathway^18^, as expressed in the temporal dynamics of cortical activity.

In this study, we extend this integrative framework by contrasting findings from human ERP experiments with results from computational simulations of the neurodynamics of self-initiated actions. This approach bridges cognitive theory, neural substrates, and experimental observations, providing a better understanding of volitional behavior.

### Objectives and goals

In light of the question posed at the beginning of the paper—namely, whether volition is instantiated by specific dynamical neural signatures that depend on the source of input, internal or external—the central focus here is to identify neurodynamic markers that differentiate these modes of action.

To tackle this issue, we propose the following hypothesis, examining perceptual decision-making processes by comparing self-initiated and externally triggered actions in terms of their neural dynamics and adaptive mechanisms.

#### Hypothesis

Self-initiated actions are preceded by distinctive neurodynamic signatures that reflect a preparatory intention process and reliably differentiate them from externally triggered actions.

This hypothesis concerns the role of underlying control mechanisms as precursors of volitional decisions and the emergence of intention leading to action across contexts that stimulate behavior. To investigate this hypothesis, we examine behavioral changes in oscillatory activity, as well as control switching reflected in the output of a neurocomputational simulation of corresponding cortical activity. As a first step, we extend the neurocomputational model developed in prior work^19^ to explore the neural mechanisms underlying externally triggered volitional actions. Subsequently, we focus on the qualitative comparison of the observations from the EEG/ERP data collected by Khalighinejad et al.^20^ with the patterns generated by our extended model, describing the similarities and differences with real EEG/ERP data in relation to the properties of our model. By comparing the simulated data with empirical EEG data, we seek to test whether empirical evidence supports the model’s prediction that self-initiated actions involve neurodynamic markers that distinguish them from externally triggered movements. By comparing the simulated data with the EEG real data, we aimed to test whether empirical evidence supports the model’s prediction that self-initiated actions involve a neurodynamical marker that distinguishes them from externally triggered movements.

## Methodology

### The neurocomputational model

The starting point is a previously developed neurocomputational model^19^. This construct models the emergence of self-initiated actions from the interaction between internal intentions and external stimuli, and it proposes theoretical mechanisms of volitional control in the anterior cingulate cortex (ACC) and lateral prefrontal cortex (LPFC).

The fundamental premise of the model is that endogenous signals including interoceptive states, emotional inputs, and internally generated goals are crucial drivers of voluntary action. These signals regulate motivational and contextual information in the ACC, understood as a hub transferring afferent signals from emotion-related neural structures and mesolimbic dopamine to the LPFC. In this scheme, the ACC controls the LPFC neurodynamics through direct and indirect pathways, via feedforward and feedback loops.

The ACC interacts with three LPFC subregions through Brodmann areas (BA) 9, 10, and 46, facilitating hierarchical control through reciprocal connections. The ACC projects to BA9, where attentional filtering and orienting processes integrate endogenous inputs with information from the external environment. This integration in BA9 contributes to the activation of long-term stored goals and their associated action plans. Signals from BA9 are further transmitted to BA10, which is involved in regulating goals and mediating decisions about which internally generated intentions should be prioritized. BA10, in turn, communicates with BA46, where more concrete action representations are encoded and where processes such as evaluating action-outcome contingencies and predicting the consequences of different behavioral choices occur^21,22,23,24,25,26,27^. Thus, our model outlines a hierarchical decision-making process involving specific LPFC subregions that contribute to different levels of volitional control. It suggests that final decisions follow a structured sequence, in which the control of stored goals guides the regulation of associated actions according to the evaluation of outcomes.

The model predicts that self-initiated actions are accompanied by a shift in neural dynamics. Specifically, theoretical neural systems transition from a chaotic, high-dimensional state to more stable, low-dimensional attractors as an intention becomes consolidated and moves closer to execution. These changes can be evidenced by estimating the dimensionality of the attractor applying time-delayed embedding for phase space reconstruction^28^.

Also, as the attractor evolves from a chaos to stability, we observe a change in the frequency characteristics of neural signals, which is interpreted as the stabilization of internal representations necessary for intentional control. This motivated the use of similar transitions in frequency as a proxy for the chaotic-to-stable dynamical transitions, at least within the scope of the computational experiments performed here. Fig 1 provides an overview of this theoretical framework, illustrating the flow of information among cortical areas involved in volitional action.

**Fig 1.**
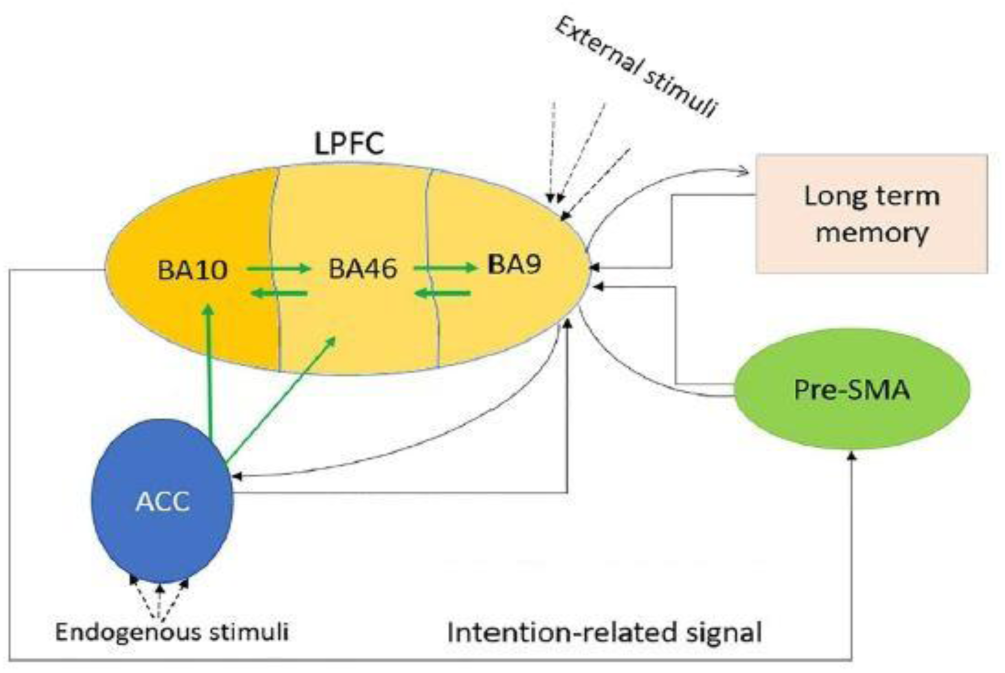
Flow of Endogenous Information Driving Self-Initiated Action. Information flow in self-initiated action shows ACC sending endogenous signals to LPFC regions (BA9, BA10, BA46), where BA9 activates goals and actions, BA10 modulates signals to guide BA46 action selection and pre-SMA preparation, while ACC encodes BA10 and BA46 activity via the dopaminergic pathway and OFC expectancy signals to direct action competition and selection (adapted from Hassannejad Nazir et al.^19^).

The comparison between the neurodynamic behavior predicted by our computational model and empirical ERP data requires additional data. First, we extend our model to produce responses for two conditions (externally-generated vs self-initiated). Then, we generate a dataset corresponding to externally-triggered actions, by simulating a scenario wherein the system transitions to alternative states beyond its current engagement in ongoing processes. This is done by adjusting the dynamical parameters that control the dynamics of the stimulating pathway and the balancing the relative influence on external vs internal stimuli.

We modified parameters related to the endogenous signal, such as ACC neural arousal (the Q value in the model) and ACC neural connection strengths, at various levels. We then analyzed the model’s behavior under different ACC input strengths. The threshold at which the system no longer behaves similarly to a self-initiated action is considered the underlying variable involved in externally-triggered actions.

Alternatively, simulating externally-triggered actions involves “stimulating” the model with significantly larger external stimuli compared to internal stimuli. To achieve this, we set variables associated with external stimuli, including frequency and amplitude, higher than those of the endogenous signal. The adjusted parameters are listed in Table 1, while all other parameters remained unchanged from the previous model^19^.

**Table 1.**
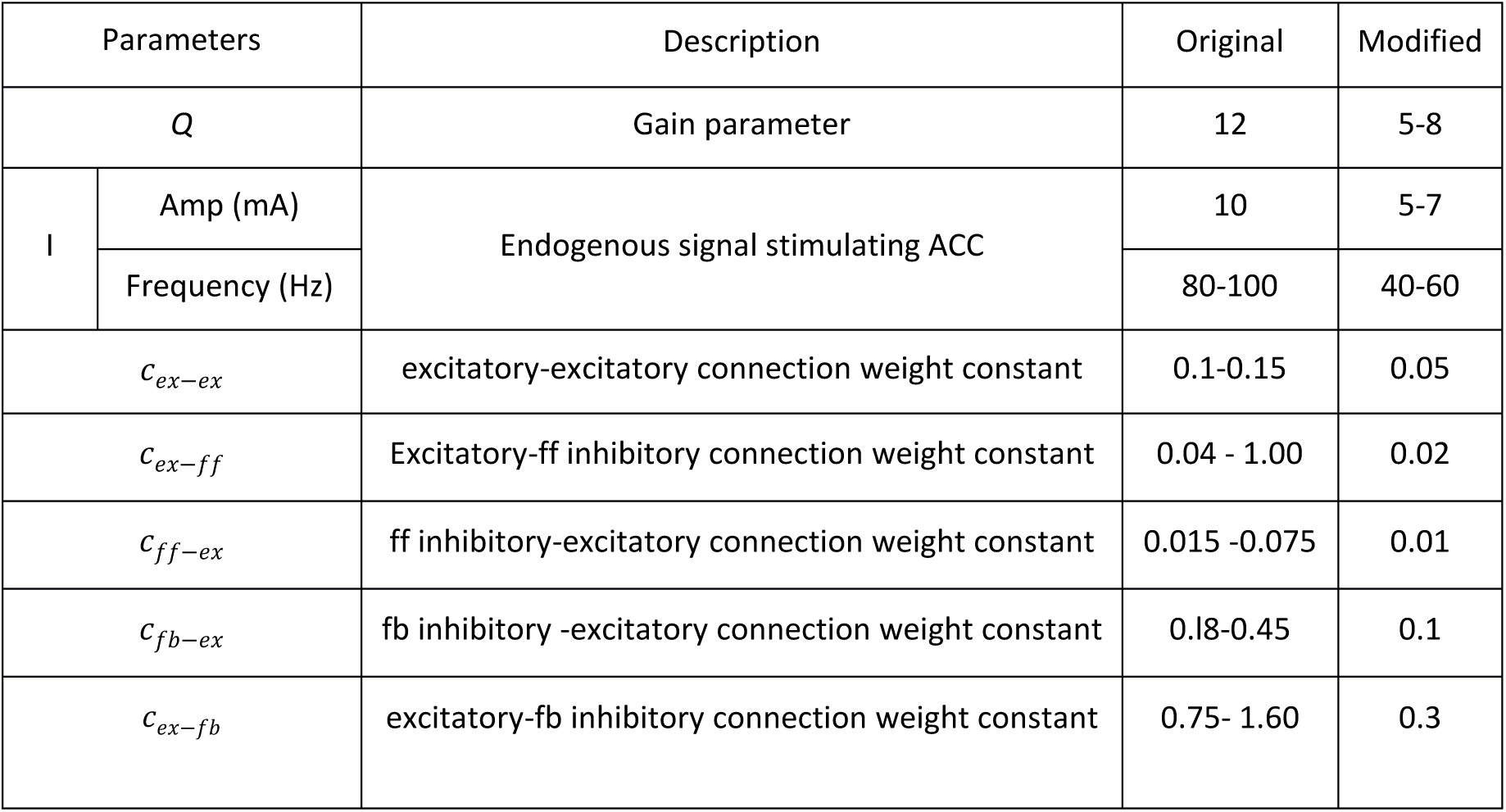
The parameters chosen for the simulation of the three-layered networks in the ACC. Many of these numbers are within physiological realistic ranges.

In continuation, we qualitatively compare the resulting predictions with ERP data collected by Khalighinejad et al.^20^ which will be described in the following section.

### A behavioural experiment

The paradigm designed by Khalighinejad and colleagues offers a clear operationalization of self-initiated actions and provides an excellent basis for comparison with our phenomenological models. As a suitable starting point for this comparison, we chose work by Khalighinejad et al.^20^ that showed differences in the readiness potentials (RPs) preceding voluntary action during a perceptual decision-making experiment, when subjects acted in response external cues vs. self-initiation.

In their paradigm, participants performed a task that allowed for endogenous “skip” responses while waiting for target stimuli. This was compared to a condition in which participants were explicitly instructed to skip, thereby balancing the frequency and timing of motor actions across conditions. The experiment operationalized self-initiated action through these voluntary skip responses in a perceptual decision task, without explicitly instructing participants to “act freely.” The inclusion of externally triggered skip trials controlled for contextual factors, thereby isolating the voluntary control component of behavior. A more detailed depiction of the experiment is provided in Fig 2.

**Fig 2.**
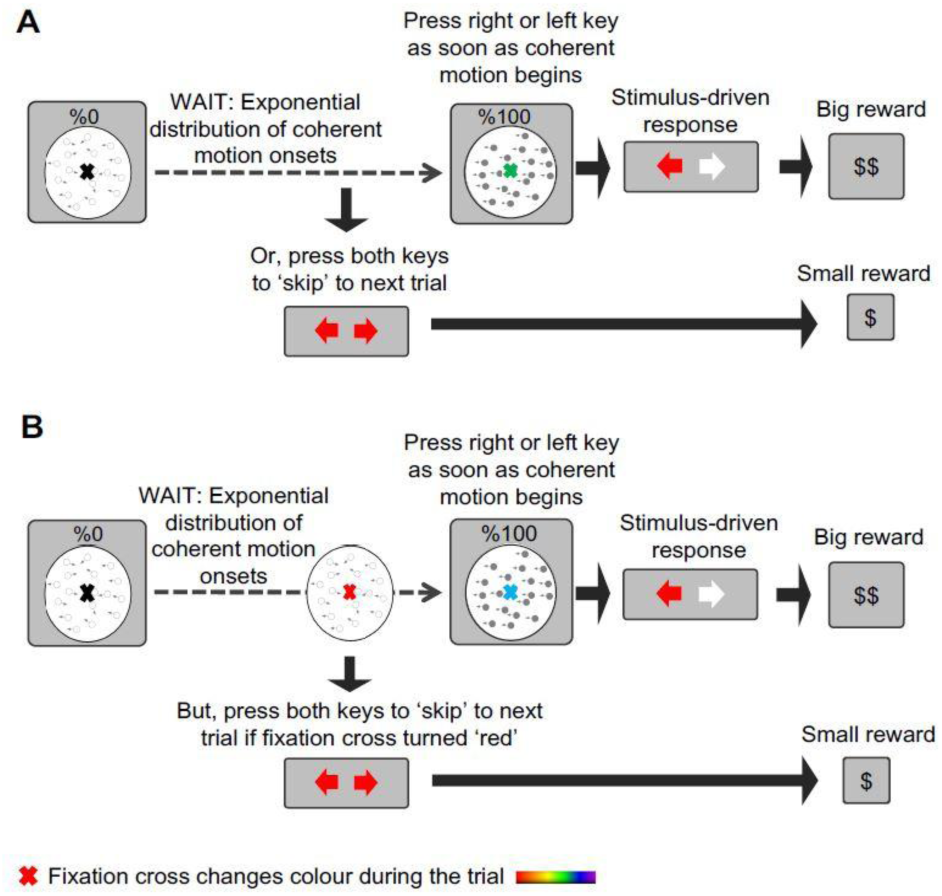
Timeline of the Self-Initiated and Externally-Triggered Decision-Making Task. This experiment illustrates the sequence of events in an experimental trial where participants responded to the direction of dot motion using left or right keypresses. The onset of dot motion was unpredictable, occurring after a delay drawn from an exponential distribution. In the ’self-initiated’ blocks, participants awaited the unpredictable dot-motion stimulus and received rewards for correctly identifying the motion direction with left or right responses. They had the option to skip long waits for the motion stimulus by performing a bilateral keypress, thus choosing between waiting, which incurred a time loss but offered a substantial reward, and ’skipping,’ which saved time but yielded smaller rewards. The color of the fixation cross changed continuously during the trial but was irrelevant to the decision task. In contrast, in the ’externally-triggered’ blocks, participants were instructed to perform bilateral skip keypresses only when the fixation cross turned red and refrained from doing so otherwise (adapted with permission from Khalighinejad et al. ^20^).

## Simulations and results

As a first step, we aim to demonstrate that our extended model can reproduce neural dynamics comparable to those observed in ERP experiments. Establishing this correspondence allows us to then hypothesize how these dynamics may reflect plausible neural mechanisms underlying volitional control. Building on this foundation, we characterize the neurodynamic signatures of preparatory states during volitional control that distinguish the processing strategies of self-initiated versus externally triggered actions in a simplified, small-scale model of cortical activity based on neural networks of limited size. This approach may help identify neural markers of intentionality in each controlling state, suggesting how the agency of each state is encoded in its dynamics.

### Network Oscillations

At the first step, simulating and averaging oscillatory activity provides a global, integrative view of how preparation unfolds across the network over time. Comparing how neural populations coordinate during these two processes may reveal differences in processing strategy between self-initiated and externally triggered actions. The observations of the model in both self-initiated and externally-triggered contexts are depicted in Fig 3. All model responses shown represent the mean across 100 simulated trials (n = 100), similar to averaging across many trials in ERP/RP recordings. This averaging provides a stable representation of network dynamics while allowing comparison with empirical EEG data.

**Fig 3.**
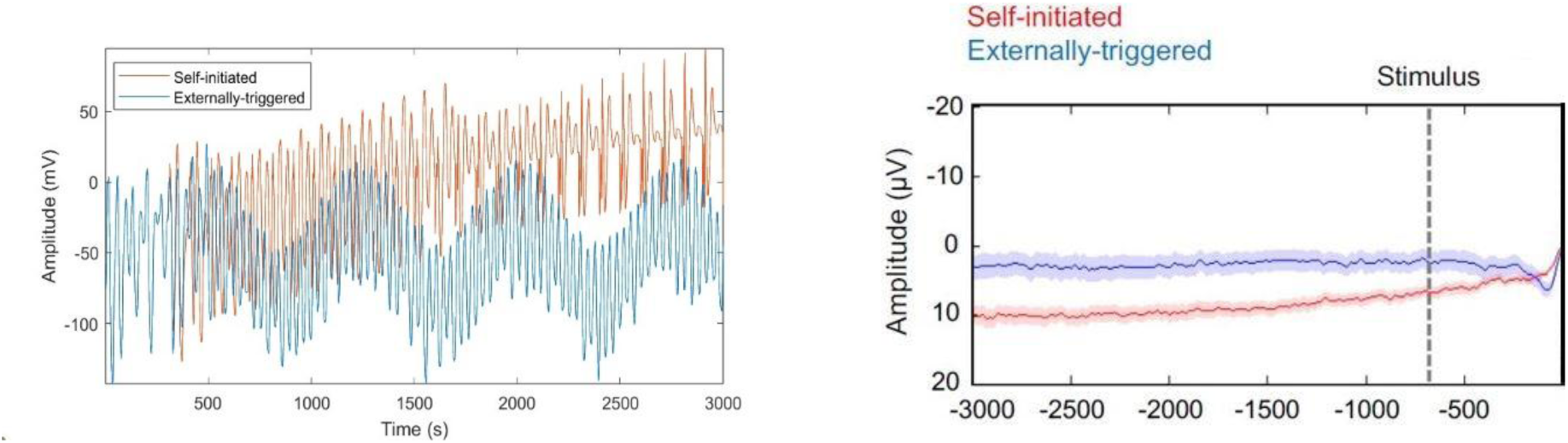
Comparison of simulated and empirical oscillatory activity preceding action. (Left) Simulation of the grand average oscillatory activity in the lateral prefrontal cortex during self-initiated (red) and externally triggered (blue) conditions. The rising trend in the self-initiated condition reflects increasing preparatory activity in BA10 and BA46, likely shaped by recurrent input from the ACC. Externally triggered actions show a flatter profile, consistent with goal retrieval initiated by external cues. Notably, both conditions display nearly identical activity before the onset of self-initiation or stimulus presentation. (Right) Empirical EEG recordings from FCz preceding skip actions in self-initiated (red) and externally triggered (blue) contexts, time-locked to action onset (vertical line). The empirical traces mirror the simulated patterns, supporting the model’s ability to reproduce key features of preparatory neural dynamics (adopted from Khalighinejad et.al.^20^).

Fig 3 illustrates the oscillatory activity of both simulated and empirical data from the control structures during the preparatory process of volitional actions. Comparing the simulated and recorded data reveals that neural population coordination involves different mechanisms depending on the type of action. In self-initiated actions, the gradual buildup of neural activity reflects the engagement of feedback loops and inhibitory regulation, producing competition among potential goals and actions during intentional preparation. Different frontal areas, such as BA10 and BA46 in the lateral prefrontal cortex guided by reinforced feedback with the ACC, exert hierarchical control over these dynamics, ultimately shaping volitional behavior.

Even during externally driven actions, individuals are exposed to a decision of whether to follow or not follow the presented stimulus. In this context, the strength of external input can produce a slight buildup of oscillatory activity. However, in the model, the applied external stimulus power—designed to make the decision fully predictable—results in largely flat oscillatory activity, reflecting the absence of internal competition and reduced engagement of feedback loops.

Having characterized the average oscillatory patterns, we now focus on variability across neural populations, which highlights how the brain maintains a landscape of competing possibilities. Measuring this moment-to-moment flexibility provides insight into how intentions emerge from a network capable of exploring, selecting, and committing to action.

### Neural Flexibility

In translating cognitive behavior into neural language, behavioral shifts—whether in the healthy brain preparing to make a decision or in a diagnosed brain under different disease states—indicate periods of high neural flexibility. Therefore, identifying variables capable of explaining and predicting actions helps capture the temporal neurodynamical behavior.

Considering that the brain’s cognitive functions inherently entail signal variability, measuring neural variability provides valuable insights into the level of neural activity across various processes^29,30^. Here, we examine the cross-trial variability of the simulated data to assess the variability of neural units during the intentional preparatory process for both self-initiated and externally triggered actions, as illustrated in Fig 4.

**Fig 4.**
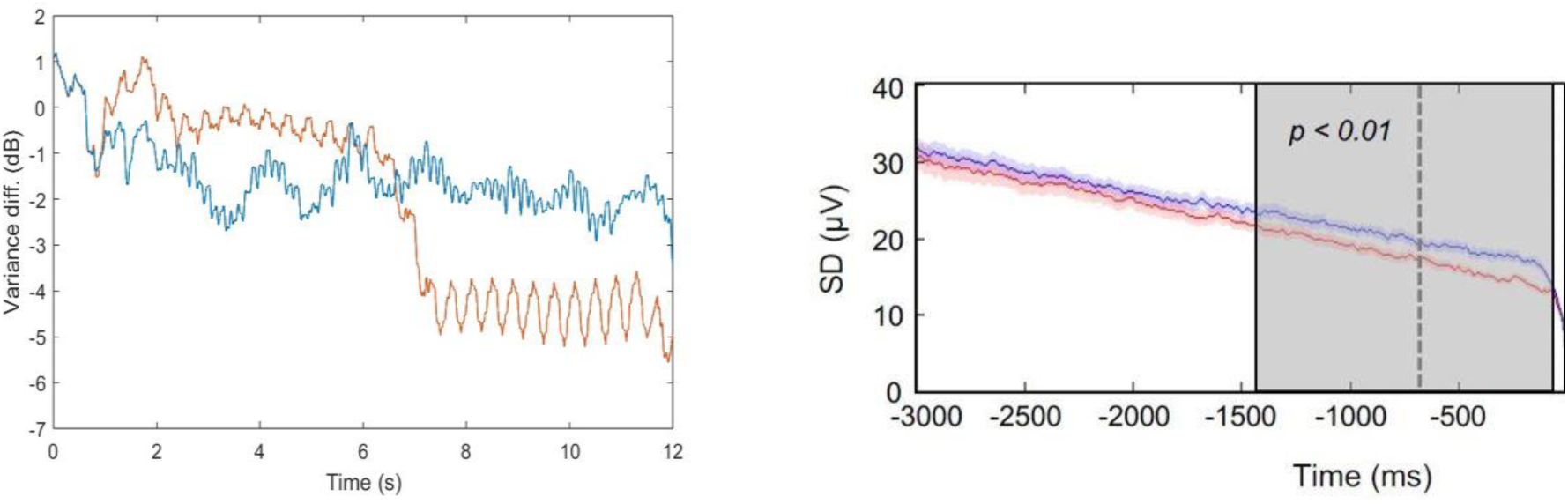
Variability of neural activity preceding action: simulated vs. empirical data. *(Left)* Baseline-corrected, log-scaled moving variance of the simulated signal during self-initiated (red) and externally triggered (blue) conditions. Variability declines more strongly in the self-initiated condition, reflecting greater stabilization of preparatory dynamics as intention consolidates. In contrast, variability remains relatively higher in externally triggered actions, consistent with a more direct, cue-driven process. *(Right)* Empirical EEG data recorded from FCz showing the trial-by-trial standard deviation of the signal preceding self-initiated (red) and externally triggered (blue) actions (adopted from Khalighinejad et al.^20^). The reduction in variability for self-initiated actions mirrors the simulated pattern, supporting the model’s account of volitional preparation as a dynamic narrowing of neural states. Action onset in both figures is indicated at *t* = 0.

The time-dependent variance of the model output during both self-initiated and externally-triggered actions in the simulated data is baseline-corrected by subtracting the mean value of the signal from - 3 s to 2.5 s, and log-scaled by conversion to decibels. The dynamic variability of the data over time provides a comprehensive insight into the fluctuations and trends within both datasets before the onset of volitional action. As depicted, both datasets show a decrease in variability, with the reduction observed in the self-initiated data being significantly greater than that observed before externally-triggered actions. To compare the variability throughout the preparatory process, the Two-sample F-test is performed for six 500 ms intervals: [-3, -2.5 s], [-2.5, -2 s], [-2, -1.5 s], [-1.5, -1 s], [-1, -0.5 s], and [-0.5, 0 s]. The test result for the interval [-1.5, -1 s] is h = 1, indicating the rejection of the null hypothesis at the default 5% significance level. The measured p-value is very small, raising doubts about the validity of the null hypothesis regarding equal variability before and after the change point in this period.

### Mean Signal Changes

Studying neural variability offers a way to understand how internal and external triggers shape the emergence of intention. This perspective highlights discrete transitions in neural dynamics as potential decision thresholds, offering a way to examine whether volitional behavior emerges from dynamic shifts during the control process. To examine these transitions, we apply change-point detection to identify moments when neural activity shifts into a new state. Fig 5 illustrates the sudden shifts in signal mean during both volitional contexts. The change points in each context mark the instances where variability undergoes abrupt alterations, potentially indicating changes in the dynamic state. The dynamic state of neural units engaged in self-initiated action undergoes a change approximately 2 seconds before the completion of the preparatory process and the onset of action. This marks the moment when signals propagate from BA10 to BA46 during the action control process, transitioning from intentional goal control to intentional action control. Conversely, change points during externally-triggered actions are noticeable early in the intentional goal control phase, with neural activity variability remaining relatively constant for the remainder of the preparatory process before action onset.

**Fig 5.**
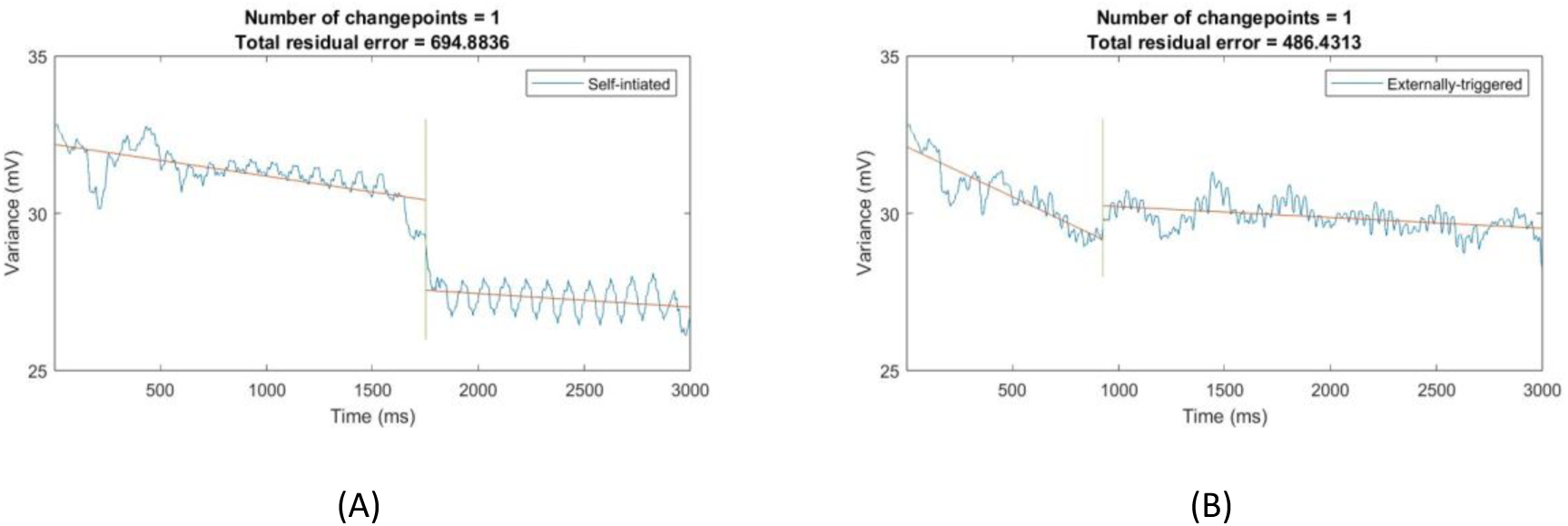
Illustration of the abrupt change in the signal mean of simulated data. A notable shift in the mean of self-initiation is observed 1247 ms before the onset of the self-initiated action, coinciding with the signal propagation from BA10 to BA46 during the intentional control process. Similarly, a significant change in the mean signal for externally-triggered actions occurs 2072 ms before the onset of the externally-triggered action during the preparatory process controlled by BA10. The change points in both subfigures represent the transition during the intentional preparatory controlling process preceding (A) self-initiation and (B) externally-triggered action. The action occurs at t = 0.

To corroborate our observation, we investigate the dynamic behavior of neural units in the frequency domain and compare it with EEG data recorded from the FCz electrode.

### Time–Frequency Dynamics

In tracing the preparatory process of intention, each analytical step adds a complementary perspective on the underlying dynamics. To capture the preparatory process of intention, it is necessary to follow how neural dynamics evolve across different dimensions. A first glimpse comes from oscillatory activity, which reflects the rhythmic coordination of neural populations. Yet rhythms alone do not tell the whole story: their stability over time also matters, and examining variability reveals whether the system remains flexible or begins to converge on a single outcome. At certain points this convergence becomes decisive, marked by abrupt transitions in the signal that signal a shift from preparation toward execution. These stages together set the stage for the final perspective, here. Frequency analysis integrates these perspectives by showing how oscillatory content itself evolves during this process. Because EEG signals are commonly characterized in the time–frequency domain, this last step provides both a unifying view of the preceding dynamics and a direct point of comparison with empirical data.

As Fig 6 illustrates, the frequency content of simulated signals during self-initiation and externally-triggered processes is examined using spectrograms. A gradual transition in frequency content from beta to gamma during the intentional preparatory process of self-initiated actions can be observed. In contrast, the frequency content remains largely constant throughout the entire intentional period of the externally-triggered context, with minimal changes detected.

**Fig 6.**
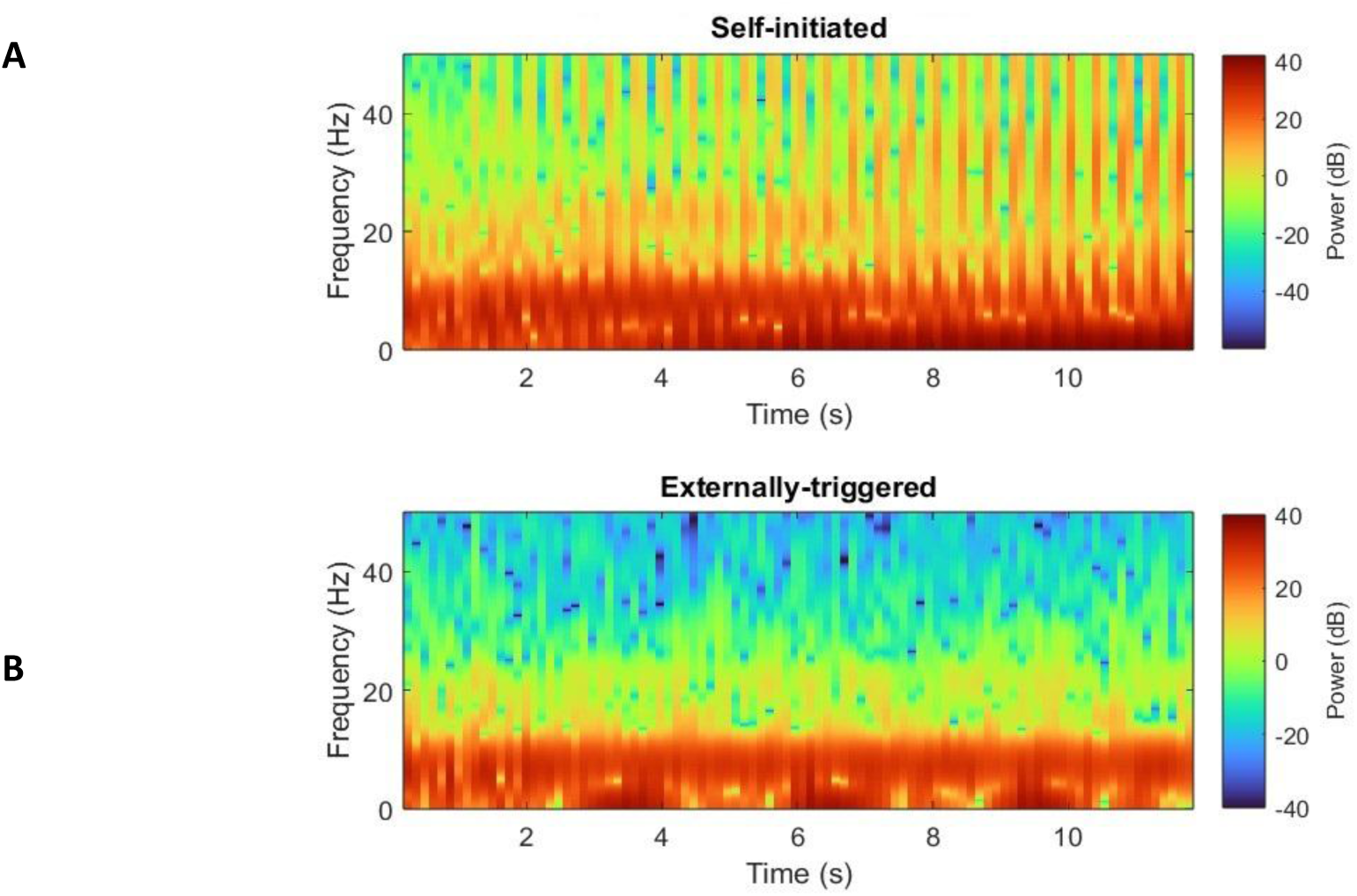
Spectral Dynamics of Self-Initiated vs. Externally-Triggered Preparatory Activity. This spectrogram illustrates the frequency content and time-varying characteristics of the signals relative to baseline (3–2.5 s prior to action). It showcases the simulated signals in both self-initiated (A) and externally-triggered (B) contexts in the frequency domain. In (A), the frequency content of neural activity during self-initiated intentional preparatory processes transitions to higher power levels. In contrast, (B) shows constant frequency components during the preparatory process without any observed transitions.

The real EEG data recorded from FCz is also analyzed in the time-frequency domain to examine transitions in both self-initiated and externally-triggered conditions.

As illustrated in Fig 7, the recorded activity exhibits frequency changes in the beta band during the preparatory process of self-initiated actions (Fig 7A), whereas no such changes are observed during externally triggered actions (Fig 7B). Spectral analysis of averaged signals from the FCz electrodes reveals that the increase in beta power during self-initiated actions exceeds that observed in externally triggered actions, highlighting distinct neurodynamic requirements and neural correlates associated with intentional action control.

**Fig 7.**
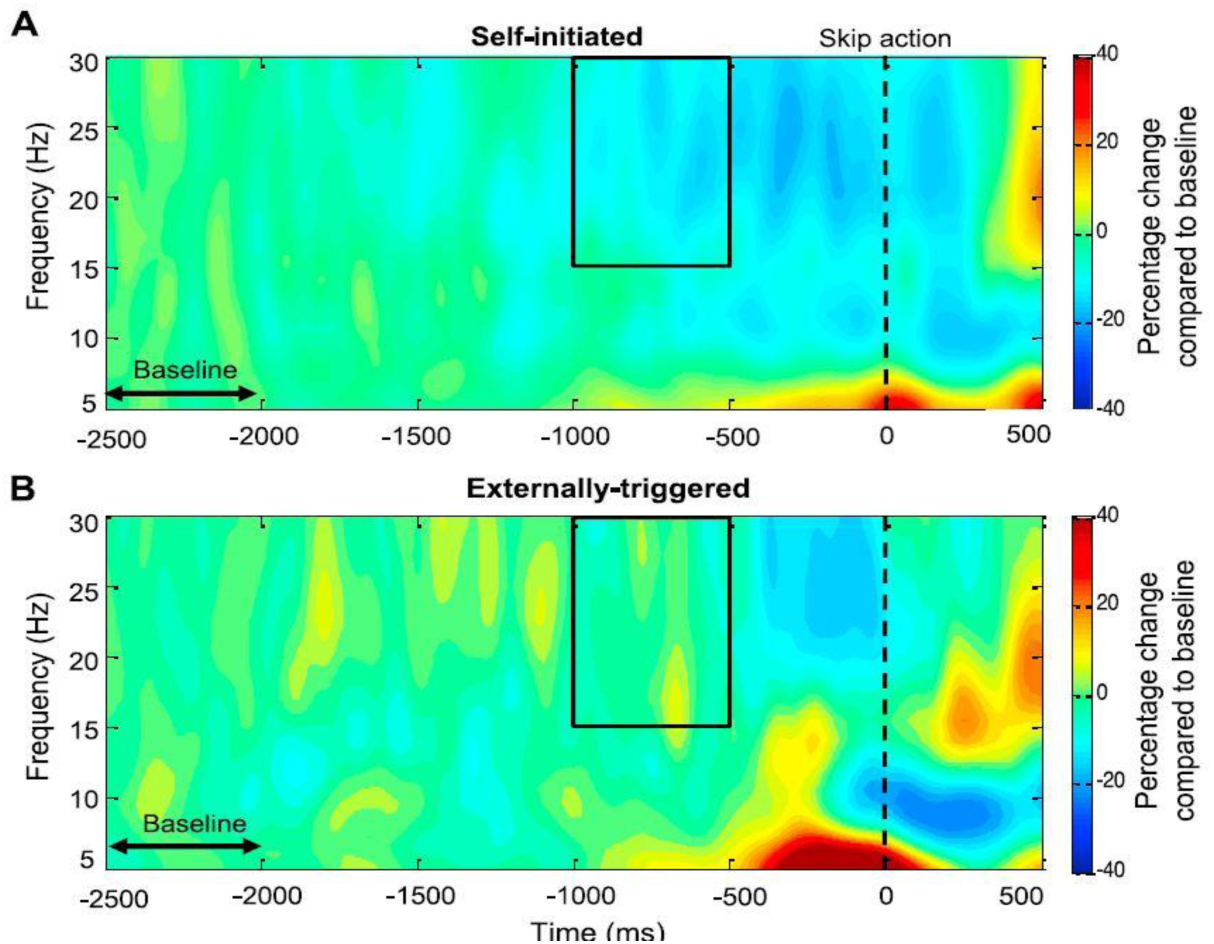
Percentage Change in EEG Power Relative to Baseline Across Action Conditions. This figure shows the percentage change in total experimental EEG power relative to baseline (2.5–2 s prior to action). (A) The percentage change in power is computed from 1 to 0.5 s prior to the skip action, focusing on the 15 to 30 Hz frequency range during the self-initiated condition. (B) Observations regarding frequency content during the externally-triggered context are presented (adopted from Khalighinejad et.al.^20^).

## Discussion

In this paper, we propose a hypothesis concerning the neurodynamics of volitional action in self-initiated and externally-triggered contexts. Our approach involves qualitatively comparing EEG observations with the results of a neurocomputational model designed to simulate cortical activity underlying these processes. The model serves as a framework for testing our hypothesis by identifying neural state transitions indicative of control switching. However, to ensure the robustness of our findings, experimental EEG data is necessary to validate the model’s predictions. Employing the neurocomputational model presented by Hassannejad Nazir et al.^19^ and taking into account the properties of specific input signals (internal), we generate two datasets corresponding to distinct categories of volitional actions.

The computational models provide a basis for better understanding the brain through new interpretations of the data. They facilitate insights into the potential underlying neural processes, which are closely linked to the nature of the developed models. The presented model attempts to describe the underlying mechanism involved in self-initiated action phenomenologically. Additionally, assessing the relationship between the gained results and the associated properties of the model provides potential explanatory insight into this field. Likewise, the variety of phenomenological approaches describing the EEG recordings broadens our knowledge about the brain. However, existing limiting factors prevent a deep exploration of the involved neural mechanisms. Therefore, qualitative comparisons between the neurocomputational findings and EEG data observations not only provide descriptive insights but also have the potential to enhance the explanatory power of our understanding. For example, we can readily manipulate model parameters to alter its behavior (to simulate “injuries”, deficits or excesses in various inputs, create conditions that resemble brain disease, etc) in the model in ways that are impossible or unethical in a human experimental setting. Hence, modeling/simulation is a convenient way to generate exploratory hypothesis that complement experimental work.

The EEG data in the perception task is recorded by N. Khalighinejad and colleagues^20^ in two conditions: self-initiated and externally-triggered. We make a qualitative comparison with these data by generating simulated data in similar volitional contexts. The baseline condition here results from the reduction of endogenous stimuli projected by the ACC to different LPFC subregions. In our neurocomputational model, the self-initiated process involves significant exchanges of influences among regions, contrasting with the externally-triggered baseline condition where some regions are inactive or involved at minimal activity levels due to the reduced excitability of the received endogenous input. This difference in neural activation is biologically inspired with brain regions that receive stronger stimulation exhibit greater responsiveness and activity, illustrated during self-initiated behavior. Conversely, when the intensity of stimulating input is decreased, these areas become dormant, exhibiting minimal activity until they become more active during self-initiated action.

As mentioned, the differences between self-initiated and externally-triggered actions lie in the source of stimulation driving the action. Therefore, the baseline conditions representing externally-triggered actions are generated by tweaking the system’s inputs (see Table I in Methodology).

In both control conditions—endogenous and externally triggered actions—we examined the neurodynamic transitions of the simulated data across both time and frequency domains during the intentional preparatory process, reflecting mechanisms of intentional control. We also compared the dynamic features of the simulated neural activity with those observed in empirical data.

### Neural Variability in the Time Domain

To examine the neural mechanisms underlying self-initiated versus externally triggered actions, we analyzed the variability of neural activity in both simulated and empirical EEG data during the intentional preparatory process. Neural variability serves as a marker for assessing the degree of involvement of a cortical region in cognitive functions, and changes in variability can reveal differences in neurodynamic organization preceding different types of actions.

Both simulated and empirical data exhibit qualitatively similar patterns of neural variability during the preparatory phase of self-initiated actions.

A decrease in variability, evident in both types of data, may indicate a state transition in both control conditions. However, the reduction observed prior to self-initiated action onset suggests a more organized and coordinated neural state compared to baseline. This reduction is markedly greater for self-initiated actions, indicating stronger regulatory control within cortical structures involved in volitional processes.

Comparing the self-initiated and externally triggered conditions highlights mechanistic differences in the preparatory phase. In self-initiated actions, the transition toward a more stable and coordinated neural state appears abrupt and is consistent across both simulated and empirical data. By contrast, externally triggered actions exhibit a more gradual or modest reduction in variability, reflecting weaker engagement of cortical control networks. These differences are consistent with the role of BA46 in orchestrating preparatory activity during self-initiated actions, whereas the weaker modulation observed in externally triggered actions may result from less pronounced interactions between ACC projections and prefrontal regions such as BA10.

Change point analysis further supports these distinctions, revealing an abrupt shift in neural dynamics during self-initiated actions, whereas the externally triggered condition lacks a comparable sharp transition.

These findings suggest that self-initiated actions are associated with distinct neurodynamic mechanisms, reflecting a higher level of cortical coordination and preparatory control compared to externally triggered actions. Overall, the results support the hypothesis that self-initiated actions involve unique neurodynamic signatures that differentiate them from externally triggered behaviors, providing insights into the neural basis of volitional control.

### Neural Synchronization and Frequency Transitions

Changes in neural variability, indicative of neural activity levels, may correlate with the synchronization of neural oscillations. Reduced synchronization can lead to increased neural variability, while enhanced synchronization brings about more stabilized neural behavior. Our observations in both simulated and empirical data suggest that the greater decrease in neural unit variability during self-initiated actions may reflect a transition towards more synchronized behavior. This transition, from low to high synchrony, may coincide with frequency shifts. Higher frequencies require more aligned neural firing and greater synchronization. To explore this, we examine frequency patterns during the preparatory process preceding both self-initiated and externally-triggered actions as well.

As illustrated in Figs 6A and 7A, neural activity demonstrates frequency transitions from lower to higher frequencies before the onset of self-initiated action, signifying neurodynamic change. In contrast, the neural oscillatory activity shown in Figs 6B and 7B demonstrates no or minimal transition before the onset of externally-triggered action. Considering the observed similarities between simulated and empirical data, we hypothesize that the rise in signal power from weak beta to gamma frequency might be linked to the neural mechanisms governing neural competition and inhibitory regulation, as delineated in the neurocomputational model. Therefore, we predict that frequency changes might result from the suppression or enhancement of conflicting attractors associated with goals and potential actions. In this process, competition between various long-term stored goals and actions might contribute to the emergence of the final decision. The cognitively demanding intentional preparatory process, involving attentional control and resolution of conflicting goals, modulates prefrontal neurodynamics. However, this notion may be challenged when an individual is subjected to external triggers leading to instructed action. In such cases, minimal competition and inhibitory control processes are expected to generate the final action due to the absence of diverse goals and potential actions. The observed steady-state oscillatory activity before the onset of an externally-triggered action may indicate the linear retrieval of goals and actions associated with the observed cue.

The underlying process can be better understood by delving into the impact of the neural properties inherent in the developed model. The activity level of the engaged neural areas in the model is a composite measure influenced by factors such as the size of neural patterns, neural excitability, network intensity, strength of inhibition, and input strength projecting to specific areas. Decreasing the magnitude of each of these variables diminishes the network’s activity level. Consequently, the reduced strength of the endogenous signal, compared to the external trigger, necessitates varying levels of activity in processing the received signals.

As mentioned, the simulated data originate from a phenomenological model of the LPFC (Hassannejad Nazir et al., 2023), while the real EEG data is collected from the FCz electrode (Khalighinejad et al., 2018). The comparison of these disparate datasets and the ensuing interpretations can be attributed to the strong bidirectional connection between the real LPFC and the pre-SMA via direct and indirect pathways in the brain. Thus, the insights gained from the computational model might hold promise for predicting behavior in the pre-SMA during the intentional preparatory process.

In summary, after qualitatively comparing simulated and empirical data, we suggest that a neurodynamic transition from an irregular to an ordered state could serve as the hallmark of the intentional preparatory process preceding self-initiated actions. The observed frequency shift and reduced variability support this assertion. Conversely, no or minimal neurodynamic changes are anticipated when actions are externally triggered. Furthermore, the suggested neural mechanism and associated neural properties may help explain our observations in both empirical and simulated data.

## Conclusions

Getting back to the questions initially raised in the paper regarding the feeling of agency and the experience of controlling versus being controlled, in this study we compared simulated neural dynamics with empirical EEG data to investigate the preparatory processes underlying self-initiated and externally triggered actions. The close match between simulations and recordings supports the model and, more importantly, provides insight into the mechanisms that may govern volitional control in the brain.

The results suggest that internally generated (endogenous) signals, unlike externally driven cues, can engage competitive mechanisms and feedback loops within prefrontal–cingulate networks, thereby driving the controlling process. In self-initiated actions, the observed neurodynamic patterns reveal alternating phases of stability and instability during preparation, which can be interpreted as neural signatures marking the approach to a decision threshold.

Externally triggered actions, by contrast, depend on the relative strength of external inputs compared to ongoing endogenous activity, with preparatory dynamics emerging only when external signals are weaker than internal ones.

Taken together, these findings illustrate a distinction between self-initiated actions, which involve intrinsic competition and gradual stabilization, and externally triggered actions, which rely more directly on the balance between external and internal signals. This distinction aligns with the idea that self-initiated actions involve more intrinsic competition, whereas externally cued actions reflect the interplay between environmental inputs and internal signals.

By comparing simulated and empirical dynamics, this work highlights the potential of simple computational models to reproduce behavioral patterns observed experimentally and to generate testable hypotheses regarding the factors shaping volitional control and decision thresholds in the human brain. While these results do not establish causal mechanisms, they suggest hypotheses about how different sources of input may shape volitional control and decision thresholds. More broadly, the present work demonstrates the potential of relatively simple computational models—despite their abstraction from the full complexity of biological systems—to reproduce rich behavioral dynamics observed in experiments. Such models offer a framework for generating testable predictions and exploring conditions that may be difficult or impossible to examine empirically, such as simulating neural disruptions associated with stroke, Parkinson’s disease, or other perturbations.

### Limitations of this approach

Computational models often face inherent limitations, as illustrated here through the simplifications and assumptions made during modeling. The neurocomputational models presented are based on neurally inspired, phenomenological networks simulating approximately 1,000 neural units within cortical structures. Abstraction and simplification are fundamental characteristics of these models, whereas EEG/ERP responses recorded experimentally reflect the aggregate activity of millions of neurons over vastly larger tissue volumes. These simplifications limit the direct biological fidelity of the models but simultaneously provide a controlled framework in which essential dynamical features can be examined. Despite their reduced scale, such models can reproduce patterns reminiscent of experimentally recorded EEG signals, enabling researchers to generate testable hypotheses and explore conditions that may be difficult or impossible to manipulate in vivo, such as simulating network perturbations or disease states. Recognizing these limitations is crucial for correctly interpreting the scope and applicability of the models while appreciating their potential to complement experimental work.

### Future directions

The present study focused on distinguishing self-initiated from externally triggered actions by examining emerging neural dynamics. Given that the strength of external stimuli and their interaction with endogenous signals are critical determinants of conscious awareness and volitional control, future investigations could explore how unconscious behaviors arise under such conditions. In particular, exposure to immediate or highly salient external stimuli may elicit responses that are better classified as non-volitional actions.

An important next step is to examine how these findings generalize to clinical populations. In the early stages of Parkinson’s disease (PD), individuals often rely more heavily on external cues, suggesting a disruption of internal initiation mechanisms^31^. Developing a meta-model that captures transitions between self-initiated and externally triggered neural states could provide insights into how these dynamics are altered in PD, and generate testable hypotheses regarding the progression of disease-related changes in volitional control.

Another promising avenue is to investigate how medial frontal damage, such as stroke involving the anterior cingulate cortex (ACC), affects volitional behavior. Clinical studies indicate that ACC lesions can impair the initiation of self-generated movements while leaving externally cued responses relatively preserved^32^. Computational modeling provides a controlled framework to simulate such lesions, allowing examination of how the disruption of specific network nodes shifts the balance between internally generated and externally driven actions, predicts behavioral consequences, and identifies network features critical for maintaining self-initiated behavior.

In future work, the model could be further extended to incorporate more realistic network architectures, additional cortical and subcortical regions, and modulatory neurotransmitter systems. Such enhancements would enable exploration of a broader range of pathological conditions and compensatory mechanisms, offering a powerful tool to generate hypotheses that complement experimental studies and guide future investigations into the neural basis of volitional control.

## Acknowledgements

This work was carried out as part of the international project ‘Neurophilosophy of Free Will. This work was carried out as part of the international project ‘Neurophilosophy of Free Will.’ We thank colleagues and collaborators for insightful discussions and feedback that contributed to the development of this study.

